# Resting state networks of awake adolescent and adult squirrel monkeys using ultra-high field (9.4T) functional magnetic resonance imaging

**DOI:** 10.1101/2023.01.08.523000

**Authors:** Walid Yassin, Fernando B. de Moura, Sarah L. Withey, Lei Cao, Brian D. Kangas, Jack Bergman, Stephen J. Kohut

**Affiliations:** Behavioral Neuroimaging Laboratory, McLean Hospital, Belmont, MA 02478; Behavioral Biology Program, McLean Hospital, Belmont, MA 02478; McLean Imaging Center, McLean Hospital, Belmont, MA 02478; Department of Psychiatry, Harvard Medical School, Boston, MA 02478

**Keywords:** functional magnetic resonance imaging, nonhuman primate, neuroimaging, ultra-high field, resting state networks, translational neuroimaging

## Abstract

Resting state networks (RSNs) are increasingly forwarded as candidate biomarkers for neuropsychiatric disorders. Such biomarkers may provide objective measures for evaluating novel therapeutic interventions in nonhuman primates often used in translational neuroimaging research. This study aimed to characterize the RSNs of awake squirrel monkeys and compare the characteristics of those networks in adolescent and adult subjects. Twenty-seven squirrel monkeys (*n*=12 adolescents [6 male/6 female] ∼2.5 years and *n*=15 adults [7 male/8 female] ∼9.5 years) were gradually acclimated to awake scanning procedures; whole-brain fMRI images were acquired with a 9.4 Tesla scanner. Group level independent component (IC) analysis (30 ICs) with dual regression was used to detect and compare RSNs. Twenty ICs corresponding to physiologically meaningful networks representing a range of neural functions, including motor, sensory, reward (e.g., basal ganglia), and cognitive processes were identified in both adolescent and adult monkeys. Significant age-related differences between the adult and adolescent subjects (adult > adolescent) were found in two networks of interest: (1) the right upper occipital region with an OFC IC and (2) the left temporal cortex, bilateral visual areas, and cerebellum with the cingulate IC. These results demonstrate that squirrel monkey RSNs are stable and consistent with RSNs previously identified in humans, rodents, and other nonhuman primate species. These data also identify several networks in adolescence that are conserved and others that may change into adulthood.

**Significance Statement:** Functional magnetic resonance imaging procedures have revealed important information about how the brain is modified by experimental manipulations, disease states, and aging throughout the lifespan. Preclinical neuroimaging, especially in nonhuman primates, has become a frequently used means to answer targeted questions related to brain resting-state functional connectivity. The present study characterized resting state networks (RSNs) in adult and adolescent squirrel monkeys; twenty RSNs corresponding to networks representing a range of neural functions were identified. The RSNs identified here can be utilized in future studies examining the effects of experimental manipulations on brain connectivity in squirrel monkeys. These data also may be useful for comparative analysis with other primate species to provide an evolutionary perspective for understanding brain function and organization.

## Introduction

Resting state fMRI (rsfMRI) is a technique used to map brain regional interactions inferred from the degree of temporal correlations between spontaneous low-frequency fluctuations (LFFs) in blood oxygen level dependent (BOLD) signal occurring in the absence of an explicit task (Friston, 1994; Biswal et al., 1995). rsfMRI can provide valuable information about the brain in a relatively short scan session and, unlike other functional neuroimaging techniques, does not require sustained attention or task performance. Investigations using rsfMRI have revealed multiple large-scale brain functional resting state networks (RSNs; Biswal et al., 1995; 2012) that are thought to subserve a variety of behavioral and cognitive domains. A hallmark of this approach is the high reproducibility of RSNs both within and between subjects—whether using seed-based (Vincent et al., 2007; Margulies et al., 2009; Babapoor-Farrokhran et al., 2013) or independent component analyses (ICA; Moeller et al., 2009, Damoiseaux et al. 2006; Pendse et al., 2011). The stability and reproducibility of RSNs thus merit their use as candidate biomarkers for neurological and psychiatric disorders and, in fact, alterations in RSNs have been identified in variety of psychiatric conditions (Bluhm et al., 2007; Auer 2008; Greicius et al., 2007; 2008; Yamada et al., 2017; Canario et al., 2021).

Many of the core functional networks that have been identified in humans (Biswal et al., 1995; 2012) also have been identified in laboratory animal species (Smith et al., 2009; 2013) including nonhuman primates (NHPs; Hutchison et al., 2011; Belcher et al., 2013; Yacoub et al. 2020; Liu et al., 2021) and rodents (Jonckers et al., 2011; Lu et al., 2012). As a result, preclinical neuroimaging has become a frequently used means to answer targeted questions related to RSNs. NHPs, in particular, are an invaluable asset to this type of investigation. In addition to their genetic, phylogenetic, pharmacokinetic, and neurobiological similarity to humans (Hutchison and Everling, 2012; Weerts et al., 2013; Royo et al, 2021), NHPs with known pharmacological and behavioral histories can be studied longitudinally under carefully controlled conditions that are rarely, if ever, possible in humans.

To date, the majority of preclinical fMRI investigations have involved macaques, which have a long history of serving as subjects in behavioral, pharmacological, and neuroscience research, or marmosets, which are small enough to serve as subjects in ultra-high field scanners that tend to have relatively small-bore size. However, squirrel monkeys provide an appealing alternative, highly accessible NHP species for neuroimaging studies: like macaques, squirrel monkeys have been widely used in many types of *in vivo* research (Abee, 1989; Royo et al., 2021) and, like marmosets, subjects are small enough (often <1 kg) for ultra-high field scanning. Importantly, when compared with marmosets, squirrel monkeys have larger brains with more pronounced gyrification (see Fig 1 in Royo et al., 2021), and their performance on cognitive tasks align more closely with that of macaques and humans than other primate species (reviewed by Royo et al, 2021; see also Williams and Glasgow, 2000) highlighting the translational value of neuroscience research using squirrel monkey subjects. Despite their value and popularity in biomedical research, few MRI studies have been conducted in squirrel monkeys and, in particular, little is known regarding their RSNs.

**Figure 1.**
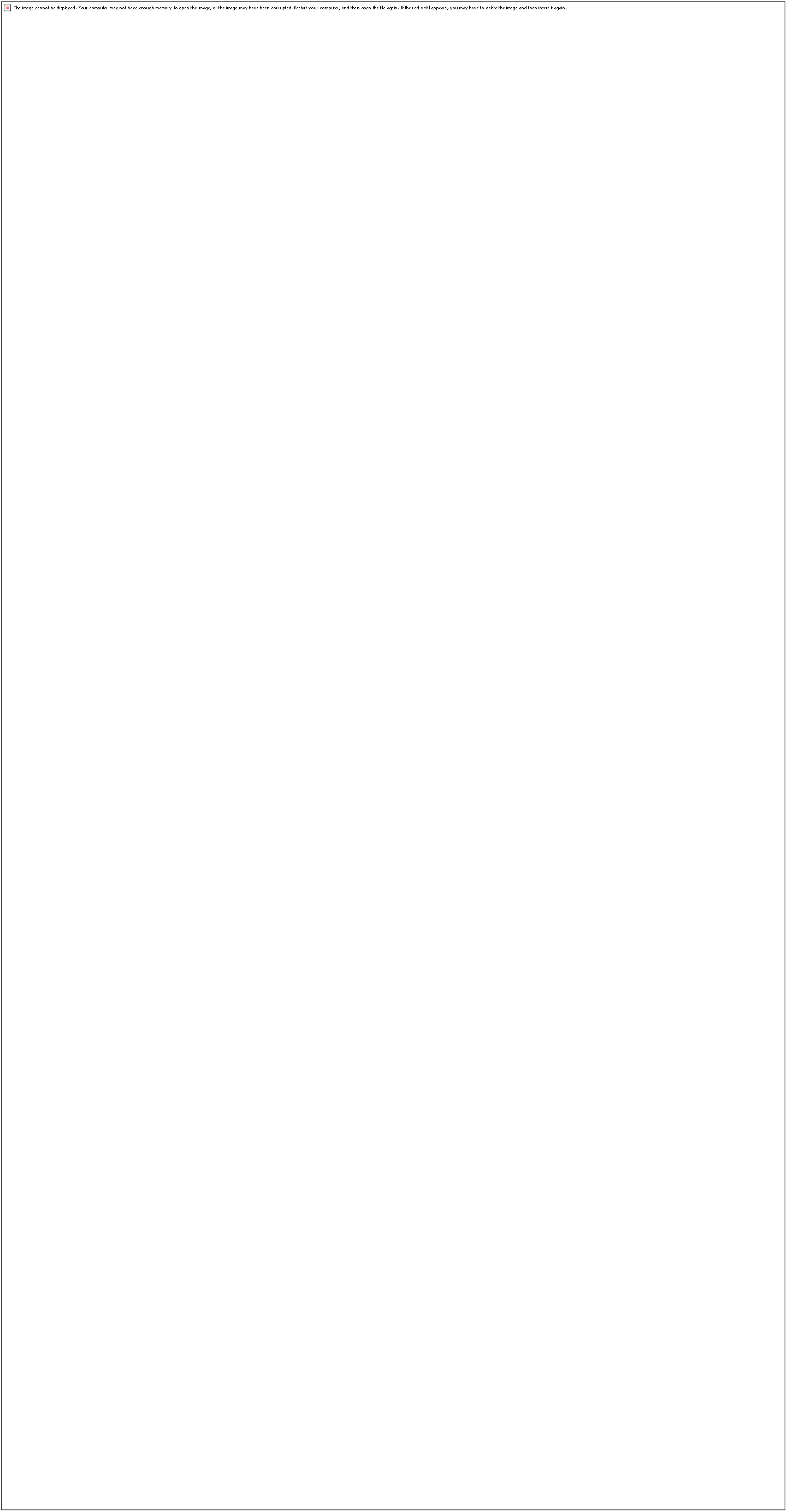
Twenty components identified as resting-state networks in the awake squirrel monkey (n = 27) using melodic group independent component analysis (ICA). Networks of similar type are presented following each other. Cingulate network (A), DMN (B), Basal ganglia (C), Thalamus (D), dmPFC (E), PFC (F), OFC (G), Salience network (H), Temporal cortex (I), Sensory motor network (J), Amygdala (K, L), Cerebellum (M), Visual network (N-S); Right visual network (N), left visual network (O), V2 and V6 visual areas (P), V1 and V2 visual areas (Q), V1, V2 and V6 visual areas (R), primary visual network (S), Temporal pole (T). Networks are shown in all three orthogonal directions overlayed on a high resolution T2w anatomical image of VALIDATE atlas. Color bar represents z-scores. ICA=Independent component analysis, DMN=Default mode network, dmPFC= Dorsomedial prefrontal cortex, PFC= prefrontal cortex, OFC= orbitofrontal cortex.

The present study sought to provide a comprehensive characterization of RSNs in experimentally naïve, awake squirrel monkeys using rsfMRI conducted at ultra-high field (9.4 Tesla). Functional neuroimaging in awake subjects avoids the potential confounding influence of anesthetic agents on functional connectivity (Lv et al., 2016; Wu et al., 2016; Zhang et al., 2019) and enhances the translational value of such data. To accomplish this, a large cohort of adolescent and adult male and female squirrel monkeys initially were acclimated to awake scanning procedures at ultra-high field. The robustness of RSNs were then evaluated in squirrel monkeys and, as well, determined whether core networks differ as a function of age, i.e., between adolescent and adult subjects.

## Materials and Methods

### Subjects

Twenty-seven squirrel monkeys (*Saimiri sciureus*) served as subjects and comprised two groups: adolescents that were approximately 2.5 years of age (mean 32.31 ±0.35 months; *n*=12; 6 male/6 female) and adults that were between 9-10 years of age (mean 117.29 ±0.74 months; *n*=15; 7 male/8 female). Subjects were housed in a temperature- and humidity-controlled vivarium with a 12-h light/dark cycle (07:00-19:00). Monkeys had unlimited access to water in the home cage and were maintained at approximate free-feeding weights with a nutritionally balanced diet of high protein chow (Purina Monkey Chow, St. Louis, MO). Fresh fruit and vitamins were provided as part of a comprehensive environmental enrichment plan. The experimental protocol was approved by the Institutional Animal Care and Use Committee at McLean Hospital in a facility licensed by the US Department of Agriculture and conducted in accordance with guidelines provided by the Committee on Care and Use of Laboratory Animals of the Institute of Laboratory Animals Resources, Commission on Life Sciences.

### Behavioral Training and Acclimation for Imaging Procedures

Extensive behavioral training was used to acclimate subjects to the MR procedures. Acclimation to the MRI apparatus typically occurred 5 days per week. Initially, subjects were trained to rest in a prone position on their haunches within a custom designed 3D printed (ABS plastic) chair enclosure for brief (5-10 min) sessions during which they were given access to intermittent presentations of approx. 0.1 ml of 30% sweetened condensed milk by syringe. The duration of acclimation training was systematically increased over several days to 30 min, after which acclimation to the helmet began. The helmet was intended to fit comfortably and securely; its design was based on squirrel monkey anatomic images collected in McLean Hospital’s 9.4 Tesla Varian system. The helmet was 3D-printed (ABS plastic) and lined with padding to limit motion and optimize comfort for the subjects. The helmet, which also included a platform to position a transmit/receive surface coil, was mounted to the chair body with plastic screws (shown in Figure S1). Once the subjects were acclimated to the helmet, the session duration was gradually extended to 60 min. The final phase of acclimation involved moving chaired subjects into a mock MRI bore housed within the laboratory. During mock scan sessions, recorded sounds from the scanner were played at decibels similar to those within the actual scanner (∼90-100 dB). Vital signs (e.g., heart rate and oxygen saturation [SPO2]; Nonin Model 7500FO, Plymouth, MN) were tracked and recorded at 5-min intervals throughout both training and MR sessions and, using live video-feeds, subjects were continuously monitored by a trained research assistant (VID-CAM-MONO-1 with SOF-842, Med-Associates, St. Albans, VT; 12M camera, MRC systems GmbH, Heidelberg, Germany).

### MRI data acquisition

Images were acquired with a 9.4 Tesla/400 mm diameter MR system (Varian Direct Drive, Varian Inc, Palo Alto, CA). The system has a 116 mm inner diameter gradient with a maximum strength of 45 G/cm; a single loop head-encompassing transmit-receive RF coil was used. Preparatory scans included automated image-based shimming <75 Hz. fMRI images were acquired with a whole-brain gradient-echo planar imaging (EPI) sequence with TE=8 ms, TR=1500 ms, and flip angle=90°; scan matrix was 64 on a 64 mm field of view with 54 × 1 mm slices; scan time for 1200 volumes was 30 min. Distortion-matched anatomic images were acquired with a spin-echo EPI sequence (TE=17.5 ms, TR=1500 ms, flip angle =90°, averages=8; scan matrix=64 on a 64 mm field of view with 54 × 1 mm coronal slices matched to the fMRI slices).

### MRI data processing

The data was visually checked slice by slice for artifacts using all three orthogonal directions. Quality control was evaluated using MRI Quality Control tool (MRIQC; Esteban et al., 2017), and group mean motion statistics were calculated for the data set. Intensity spiking was analyzed and resolved with an in-house program, “spikefix”, which is designed to find and remove spikes from fMRI datasets (https://github.com/bbfrederick/spikefix) with a threshold of 1.0 mm framewise displacement. FMRIB’s Software Library (FSL, Oxford University, UK) image preprocessing pipeline was used to process the data as follows (Smith et al., 2004; Jenkinson et al., 2012): The first 10 volumes from each scan were removed to allow for data stabilization. MCFLIRT tool (Jenkinson et al., 2002) in FSL was used for head motion correction by volume realignment to the middle volume. The resulting 12-motion correction positions were used as nuisance regressors. Session-averaged functional volumes were aligned to the VALiDATe (Schilling et al., 2017) T1w template through a 12 DOF affine transformation followed by adjustment of nonlinear distortion fields using the jip analysis toolkit (www.nitrc.org/projects/jip). Spatial smoothing was conducted using a Gaussian kernel of 2.0 mm FWHM. Registration to template space was carried out using jip. Temporal filtering was applied using a high pass filter with a 100 s cutoff (0.01 Hz).

### Independent component analysis

Group level independent component (gIC, Melodic, FSL) analysis was used to examine the resting state fMRI data. The number of independent components (ICA 30) was chosen based on previously published methods in NHPs (Lu et al., 2012; Belcher et al., 2013). The ICA method was chosen as the primary analytic strategy for this study because it has been used across a variety of species and is a highly replicable, well-validated method for investigating large-scale brain networks using rsfMRI. The widespread use and acceptance of this approach permits the ICA maps obtained here in squirrel monkeys to be directly compared to those obtained in both human and other nonhuman subjects. The robustness of the identified ICA maps was examined using exploratory analysis of 20, 25, 35, and 40 model-specified components (Belcher et al., 2013). The physiologically relevant components were visually identified and determined to be consistent with previously reported networks in humans, rhesus monkeys, marmosets, rats, and mice (Beckmann et al., 2005; Hutchison et al., 2011; Jonckers et al., 2011; Lu et al., 2012; Shirer et al., 2012; Belcher et al., 2013). The resulting component maps were overlaid on a high resolution T2w squirrel monkey brain template “VALiDATe” image. Brain regions were identified and labeled by three independent observers (WY, FBM, and SLW) and based on Royo et al. (2021) for cortical, and Gergen and MacLean (1962) for subcortical anatomy. The robustness of the networks was evaluated by visually inspecting the single subject IC networks in all subjects. The above analyses were performed for both the adolescent and adult groups, combined and separately.

### Dual regression

Subject-specific versions of the spatial maps and associated time series were generated with the spatial maps from the group average analysis using dual regression (Beckmann et al., 2009). Then, differences between the adolescent and adult groups were assessed using Randomise in FSL (two-sample unpaired t-test with covariates, using 10,000 permutations). This was run twice independently, once with motion and sex as covariates and another with motion, sex, and age. Clusters were produced using threshold-free cluster enhancement. The number of RSNs of interest was accounted for using Bonferroni correction with the original threshold set at P<0.05 (FWE) and adjusted to P<0.002 (FWE).

## Results

### Awake fMRI Data Quality

All subjects adapted well to awake MRI acclimation procedures and completed the 30-min scan session without incident. Heart rate and SPO_2_ remained stable and within normative ranges throughout the duration of each scan session. Mean motion during the scan sessions was 0.167 ± 0.04 mm for the adolescent subjects and 0.193 ± 0.02 mm for the adult subjects. No significant difference in motion was found between the two groups (P=0.6).

### Resting State Networks

All ICs representing white matter, cerebrospinal fluid, or physiological noise were excluded. This resulted in the identification of 20 ICs, out of the initial 30, corresponding to physiologically relevant networks that are consistent with published RSNs (Beckmann et al., 2005; Hutchison et al., 2011; Jonckers et al., 2011; Lu et al., 2012; Shirer et al., 2012; Belcher et al., 2013). These putatively labeled networks are shown in Figure 1, panels A-T, with additional slices of each IC shown in the supplementary material (Figures S1, sA-T), and are described as:

- Posterior cingulate network (Fig. 1A) – This network contains the bilateral posterior cingulate cortex (PCC) and the caudate nucleus.
- Posterior cingulate-parietal network (Fig. 1B) – This network includes areas that correspond to the Default Mode Network reported in humans (Abou-Elseoud et al, 2010) and contains the precuneus, PCC, and inferior parietal cortices.
- Basal ganglia network (Fig. 1C) – This network includes the head of the caudate, globus pallidus, and putamen.
- Thalamic network (Fig. 1D) – This network is localized to the thalamus with limited activation in the PCC.
- Three PFC networks were identified - The three bilateral PFC ICs contain the DMPFC (Fig. 1E, which corresponds to the anterior component of the Default Mode Network; Abou-Elseoud et al, 2010), rostral PFC (Fig. 1F) and OFC (Fig. 1G) and are named accordingly.
- Ventral-parietal network (Fig. 1H) - This network resembles the Salience Network reported in humans (Seeley et al., 2007; 2019) and includes the bilateral ventral parietal cortex, secondary somatosensory area (PV/S2), anterior parietal cortex (APC), primary motor cortex, and premotor cortex.
- Temporal network (Fig. 1I) - The temporal cortex was identified in one network localized to mid-temporal cortex and brainstem.
- Dorsal-parietal network (Fig. 1J) – This network contains brain regions associated with sensorimotor networks (cf. Rocca et al, 2009). The posterior parietal network contains the bilateral APC, bilateral posterior parietal cortex (PPC), bilateral PCC, and bilateral PV/S2.
- Amygdala-hippocampal network (Fig. 1K) – This network includes the bilateral amygdala and the hippocampus.
- Amygdala-temporal network (Fig. 1L) – The bilateral amygdala and temporal cortex are included in this network.
- Cerebellar network (Fig. 1M) – This network is limited to and covers the entire cerebellum.
- Anterior temporal network (Fig. 1T) – This network contains the bilateral temporal poles.

The occipital cortex was found in 6 of the 20 components. There were two unilateral visual components, one right component (Fig. 1N), and one left component (Fig 1O) each containing visual areas V1-4. One of the six was restricted to the bilateral primary visual area (V1 (Fig. 1S)); one contained visual areas V1 and V2 (Fig. 1Q), one contained V1, V2 and V6 (Fig. 1R) and one contained V2 and V6 (Fig. 1P).

The RSNs identified in the group level analysis described above were also found in the two groups (adults and adolescents) when analyzed separately.

As the intention of this study was not to focus on a particular network per se but to identify all the potential RSNs evident in squirrel monkeys, we did not select the model order based on whether a particular network was intact or not. However, in model order 30 the anterior and posterior components of the DMN, considered to be a key target in neurodevelopmental and neuropsychiatric disorders (see Supekar et al, 2010; Choi et al, 2021), were found in separate ICs. Lower model orders were further evaluated to determine if the intact DMN, i.e., the posterior and anterior components, could be identified. Figure S3 shows the intact DMN observed in model order 15. While the DMN was found at model order 15, several other networks were found to overlap suggesting that this model order was not optimal for a full assessment of squirrel monkey RSNs.

### Dual regression

We were further interested to know whether RSNs related to cognition are conserved between adolescent and adult subjects and, to address this, focused on networks containing cingulate, amygdala, PFC, insula, basal ganglia, and thalamus. Figure 2 shows that when motion and sex were entered as covariates, significant differences between the age groups were found in two of the networks of interest: (1) in the right occipital region (Fig 2A) with the OFC network (Fig 1G) and (2) in the left temporal cortex, bilateral occipital cortex and cerebellum (Fig 2B) with the posterior cingulate network (Fig 1A). In both networks, adults showed greater connectivity compared to adolescent subjects; P<.003. No voxels survived in any of the ICs when age was entered into the above model, even at a liberal threshold (P=.05).

**Figure 2.**
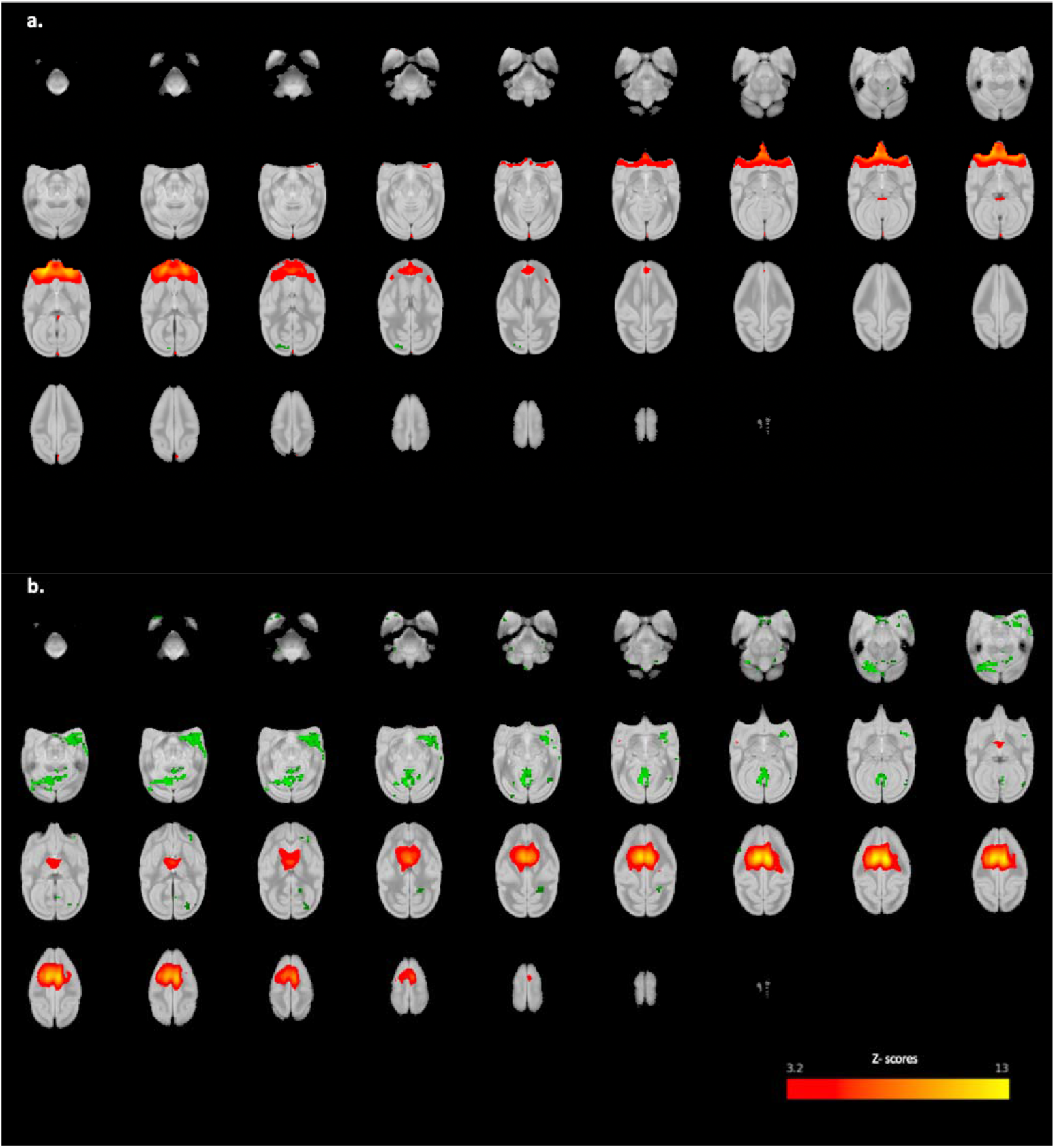
Axial brain view showing the dual regression results between adult and adolescent groups. Differences were noted in a. the OFC network in the right occipital region and b. the cingulate network in the left temporal and bilateral occipital cortices and cerebellum between the adult and adolescent subjects. The coordinates are showing the DR results in green (adult > adolescent), P <.003 overlayed on the IC network shown in Fig 1.

## Discussion

Resting-state networks are associated with key neurological and cognitive processes, and alterations of such networks are thought to reflect underlying pathology. As NHPs become increasingly utilized in neuroimaging studies (see Special Issue *Neuroimage* Vol. 244), a comprehensive mapping of functional brain networks will facilitate their use as translational models in neuropsychiatric research— both elucidating underlying mechanisms of neuropsychiatric disorders and developing targeted treatments for their remediation. With this in mind, the goal of the present study was to characterize RSNs in both adolescent and adult squirrel monkeys at ultra-high field strength (9.4 Tesla). Research using squirrel monkeys as experimental subjects has contributed importantly to our understanding of a range of psychiatric and neurodegenerative disease states (reviewed by Royo et al, 2021) and fills an important phylogenetic gap between rhesus macaques and marmosets; two NHP models used in a growing number of neuroimaging datasets. Furthermore, the present work was conducted in the absence of anesthetic agents-i.e., subjects were awake during scan sessions—and 20 networks representing a range of neural functions, including motor, sensory, reward, and cognitive processes were identified.

The RSNs described in the current report are defined by anatomic localization and classified as large-scale networks informed by anatomical features of RSNs reported in previous studies, where appropriate (Hutchinson et al., 2011; Heine et al., 2012; Belcher et al., 2013; Yacoub et al., 2020). A survey of the literature shows that the results presented here in squirrel monkeys are highly consistent with those described in a number of laboratory animals, including rodents (Bajic et al., 2017; Becerra et al., 2011; Hutchinson et al., 2010; Lu et al., 2012, Moeller et al., 2009), voles (Ortiz et al, 2018), ferrets (Zhou et al, 2016), and other NHP species (Belcher et al., 2013; Hutchinson et al., 2011). The ICs presented here contain a mixture of those that encompass both cortical and subcortical brain regions and represent a range of functions. One of the prominent features of RSNs in preclinical imaging studies is local connectivity rather than the long-range connections that are sometimes reported. For example, resting-state networks such as the DMN, which is implicated in many cognitive, social, and emotional functions (Whitfield-Gabrieli and Ford, 2012) has often been reported as being “broken” into its subcomponents; the DMN in animals has been discussed in detail by Hutchison and Everling, 2012; see also Liu et al, 2019; Garin et al, 2022. Previous studies have shown the DMN to contain both posterior and anterior components such as the PCC, precuneus, inferior parietal cortex, mPFC, and, less consistently, the hippocampal formation (Broyd et al., 2009). Although the ICs containing DMN-associated regions reported here (Figs 1E and 1B) do not include this full complex of structures, the overall pattern of connectivity identified over multiple ICs strongly suggests evidence of a squirrel monkey DMN ortholog (cf. Hutchison and Everling, 2012; Garin et al, 2022), albeit separated into regionally defined subcomponents (i.e., anterior and posterior). An assessment of various ICA decompositions of the current data yielded a DMN-like network in a lower model order (i.e., 15) that contained both anterior and posterior components (see Supplementary Figure S2). Similar partitioning of large-scale networks, including into dorsal and anterior portions of the DMN (c.f. Abou-Elseoud et al, 2010), has been reported in human fMRI data using ICA (He et al., 2009; Doucet et al., 2011). There are two likely explanations for the partitioning of larger scale networks among multiple ICs: First, ICA captures unique subnetworks that function dynamically over time, each with specific roles in neural processing, consistent with hierarchical functional organization of the brain. This may best be seen in the present study across the 6 ICs associated with visual processes that contain subregions of visual cortex (see Royo et al., 2021). Next, the distinction may have a foundation in evolutionary biology as a recent study comparing features of the DMN in humans, macaques, marmosets, and mouse lemurs found that mPFC was weakly connected to PCC in primates when compared to human (Garin et al, 2022). Regardless, the fact that each of the RSNs reported here were observed in all individual subjects emphasizes the robustness of the current data, and in turn, provides confidence that these networks also will be evident in future imaging studies with squirrel monkeys.

It is widely understood that the brain continues to develop from adolescence into adulthood (Blakemore, 2011; Rubia, 2013). The adult brain is believed to contain networks that consist of well-defined spatiotemporal connectivity, a reflection of increasing neural and hierarchal segregation and pruning as one ages (Stevens et al., 2009). Some studies have demonstrated that key RSNs-e.g., DMN, and executive network-are largely similar between children and adults; however, some studies have shown activation of frontal regions in the DMN to be weaker in children than adults, and cingulate activation was not observed in several RSNs in children (Stephens et al., 2009; Muetzel et al., 2016). In the present study, overlapping RSNs were found in both the adolescent and adult brains, which is largely consistent with findings in human subjects. However, in contrast to studies in humans, connectivity in the PFC and cingulate networks to occipital and temporal regions, respectively, was slightly greater in adult squirrel monkeys than in adolescents. While the basis for these differences is not yet known, several factors deserve consideration. First, the human literature typically includes subjects ranging in age from early to mid-adolescence for comparisons of age-related rsfMRI whereas adolescent squirrel monkeys in the present study were a relatively homogenous group, with ages spanning only a 3-month range. Thus, it is possible that the differences between adolescents or adults identified here may reflect the well-controlled age range of the subjects studied. Second, it is possible that other types of analysis (ROI-based, graph theory, dynamic connectivity analyses) or different experimental designs (task-based fMRI) may reveal other subtle differences in brain function or connectivity that are not evident using an ICA approach. While future studies are needed to answer these questions, the results of this study demonstrate the robustness and reproducibility of the ICA networks identified here in individual subjects and across different cohorts. Presumably, these findings can serve as a platform for future studies to investigate other variables or influences on rs-functional connectivity.

Patterns of functional activation in response to various stimuli and/or treatment conditions in previous studies in squirrel monkeys (Nelson et al., 2006; Gao et al., 2014; Schilling et al., 2017; Wu et al., 2017) generally have mirrored findings from the human literature (Heine et al., 2012). For example, presentation of a noxious thermal stimulus to squirrel monkey during ultra-high field imaging resulted in fMRI responses in numerous regions (e.g., thalamus, caudate, posterior insula) that are related to nociception in humans (Wu et al., 2017). Also, repeated exposure to cocaine has been shown to alter putamen and dACC connectivity (Kohut et al., 2020), a finding that recapitulates phenomena observed clinically in humans with substance use disorders (Hu et al, 2015). These results suggest the existence of orthogonal functional networks in squirrel monkeys and humans. The current report further strengthens the evidence for this idea by describing RSNs in awake squirrel monkeys, i.e., without the influence of anesthetic agents, using conditions that are analogous to those in human resting-state studies. Overall, the present findings indicate that squirrel monkeys can be used to characterize the functional neurobiological consequences of behavioral and pharmacological manipulations in highly controlled environmental and experimental conditions that are not possible in research with human subjects. Importantly, the capacity to conduct such research in awake squirrel monkeys without the influence of anesthesia further increases the translational validity of squirrel monkey research into RSNs and, more generally, brain functional connectivity.

Despite the reliability, robustness, and strength of the ICA method, it is important to acknowledge its limitations, especially sensitivity to motion (Parkes et al. 2018) requiring the removal of motion-related ICs from the data. This is typically done based on the assessor’s judgment and requires specific knowledge of the RSNs, especially in animal neuroimaging, or the use of more rigorous motion correction tools (Parkes et al. 2018). The uncertainty in determining the number of ICs in the ICA method is also important and, as mentioned earlier, may impact RSNs. Machine learning techniques may be useful for addressing this challenge (Wang and Li 2015). Future studies highlighting the connectivity patterns of particular networks of interest also might benefit from implementing a seed-based analytical approach or methods such as connectome, graph theory, or dynamic ICA analyses to delineate different hub locations and to characterize how well these hubs are connected to different brain nodes as well as the signatures of the long and short distance connections between nodes.

## Acknowledgments

R01DA048150 (SJK), R01DA047575 (JB/SJK/ BDK/SLW), Alkermes Pathways Research Awards® grant (SLW), T32DA015036 (FBM), K01DA039306 (SJK), and S10RR019356, and by the Counterdrug Technology Assessment Center, an office within the Office of National Drug Control Policy, via Contract No. DBK39-03-C-0075, awarded by the Army Contracting Agency. The content of the information does not necessarily reflect the position or the policy of the U.S. Government, and no official endorsement should be inferred.

The authors thank Jessi Stover, Samantha McGouldrick, Bryan Carlson, and Craig Stone for their efforts in acclimating subjects to awake MRI procedures, Kenroy Cayetano for engineering the experimental equipment used for these studies, Bonnie Adams for assistance with MRI data acquisition, and Drs. Dionyssis Mintzopolous and Michael L. Rohan for developing the acquisition sequences used here. We acknowledge Dr. Lisa Nickerson from the Harvard Catalyst Biostatistics consulting program for helpful discussions about independent component analysis and presentation of results. Portions of this work were presented at the annual meeting of the Society for Neuroscience, 2021.

## Author Contributions

Walid Yassin – Designed research, performed research, analyzed data, wrote the paper

Fernando B. de Moura – Performed research, analyzed data, wrote the paper

Sarah L. Withey - Performed research, analyzed data, wrote the paper

Lei Cao – Analyzed data, wrote the paper

Brian D. Kangas – Designed research, wrote the paper

Jack Bergman – Designed research, wrote the paper

Stephen J. Kohut – Designed research, performed research, analyzed data, wrote the paper

