## Supplementary Data for "Resting state networks of awake adolescent and adult squirrel monkeys using ultra-high field (9.4T) functional magnetic resonance imaging"

**Supplemental Materials**

**Figure Captions**

**Figure S1.** Schematic of the MRI apparatus developed for squirrel monkeys. Acclimation procedures are described in text.

**Figure S2.** Several axial slices of each network (sA-sT) corresponding to networks A-T in Figure 1 of and described in main text of manuscript. Same threshold level applied.

**Figure S3.** The default mode network with anterior and posterior components at a lower model order with an independent component number set at 15.

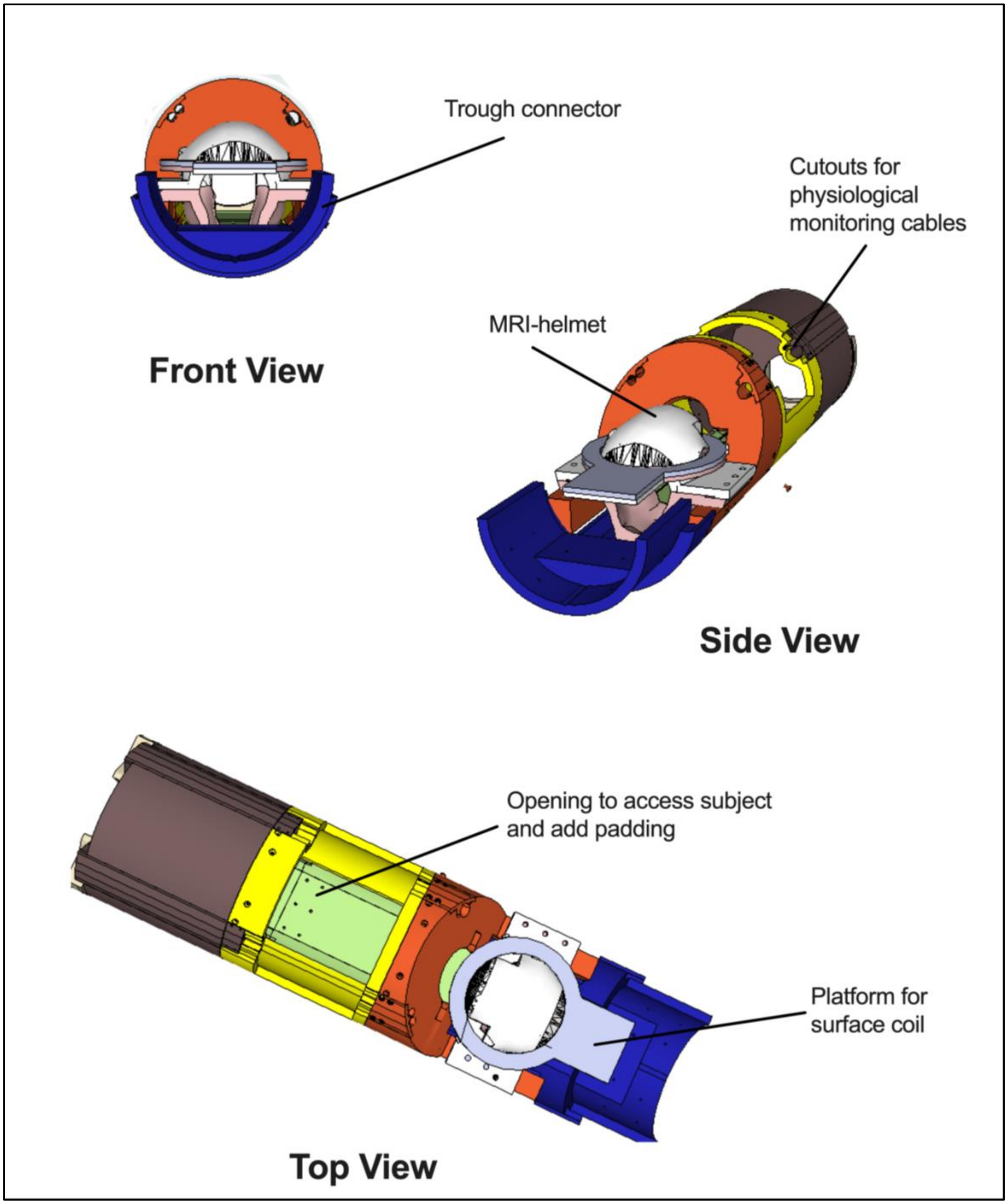

Trough connector
Front View
Cutouts for
physiological
monitoring cables
MRI-helmet
Side View
Opening to access subject
and add padding
Platform for
surface coil
Top View


**Figure S1.**

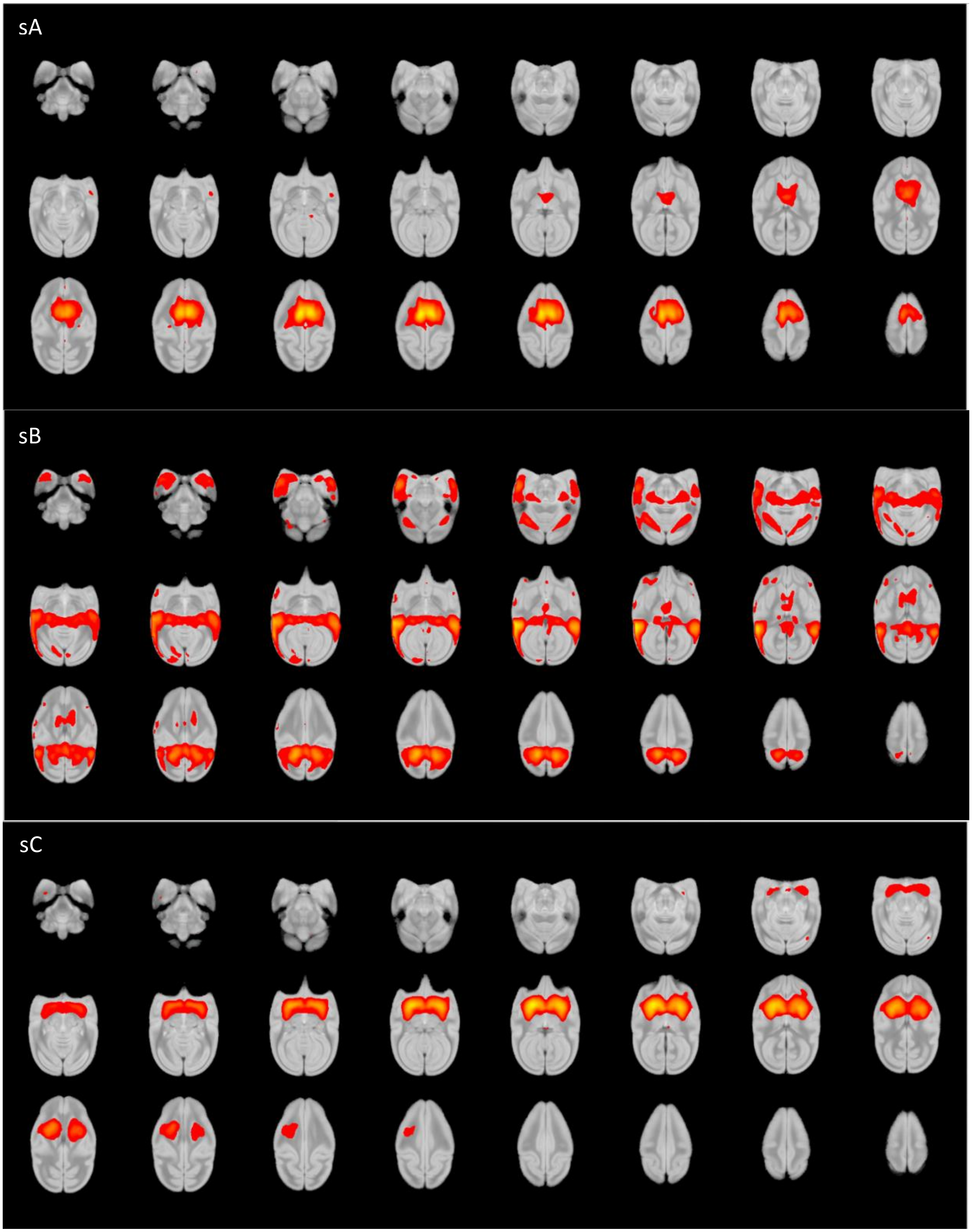
sA
sB
sC

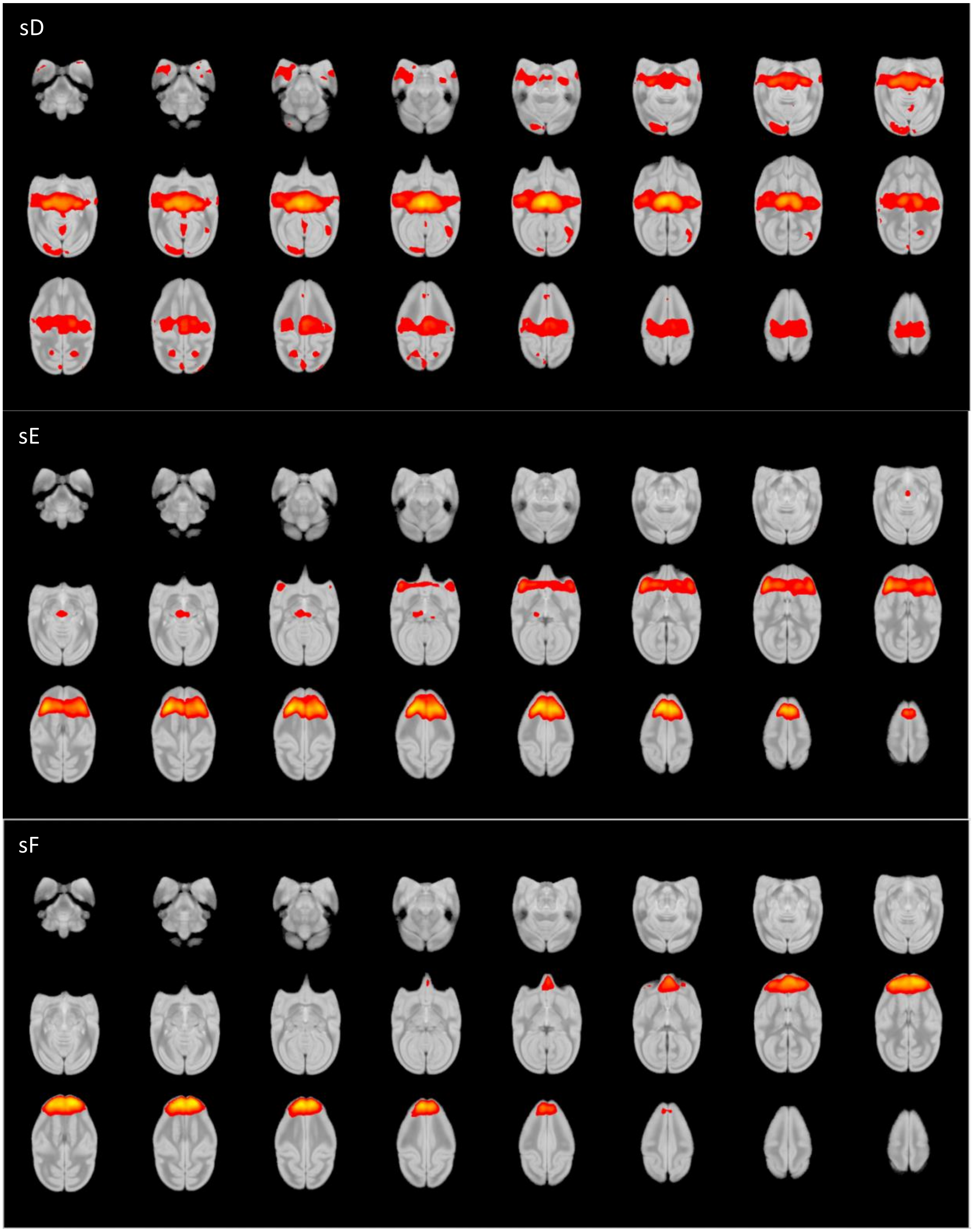
sD
sE
sF

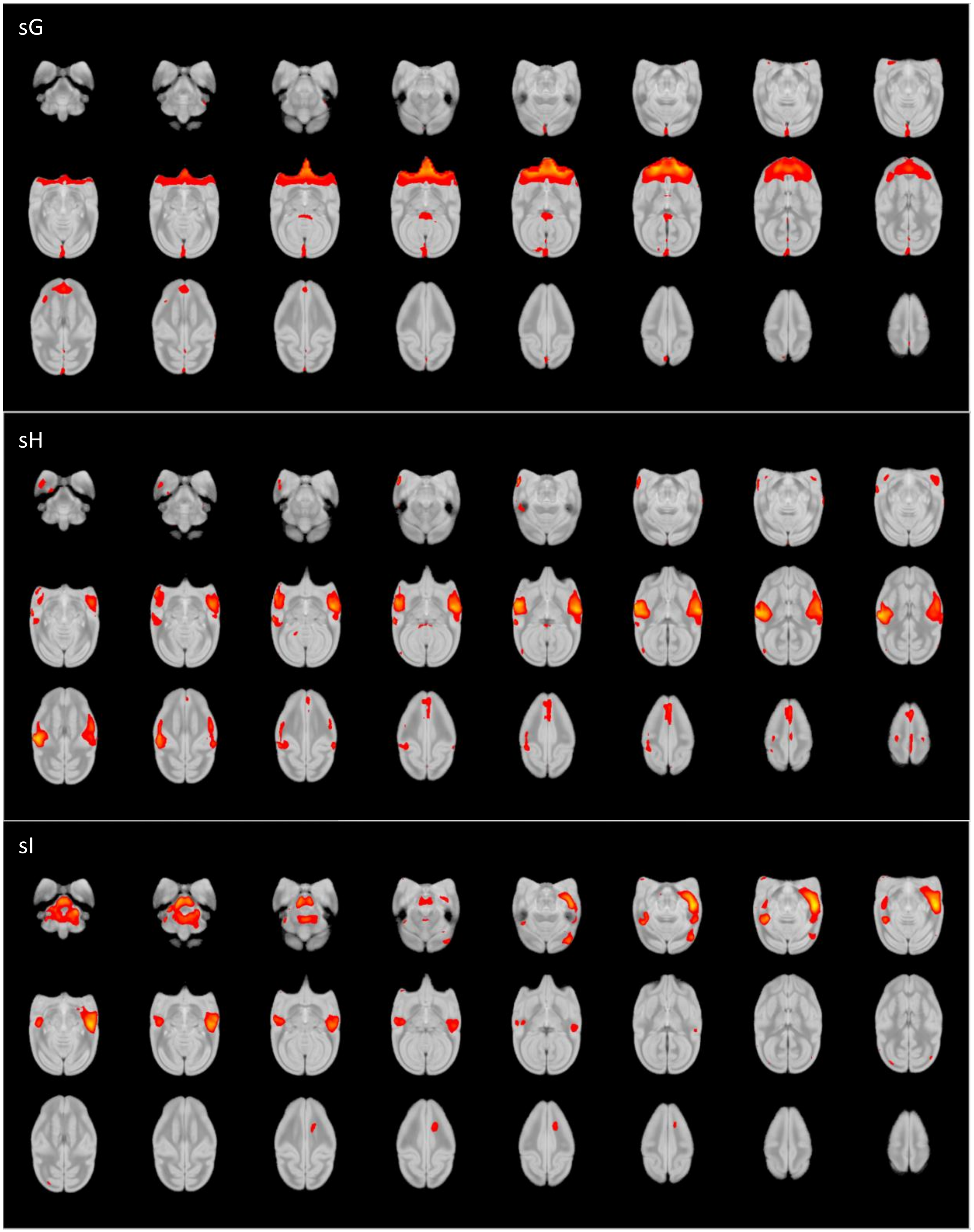
sG
sH
sI

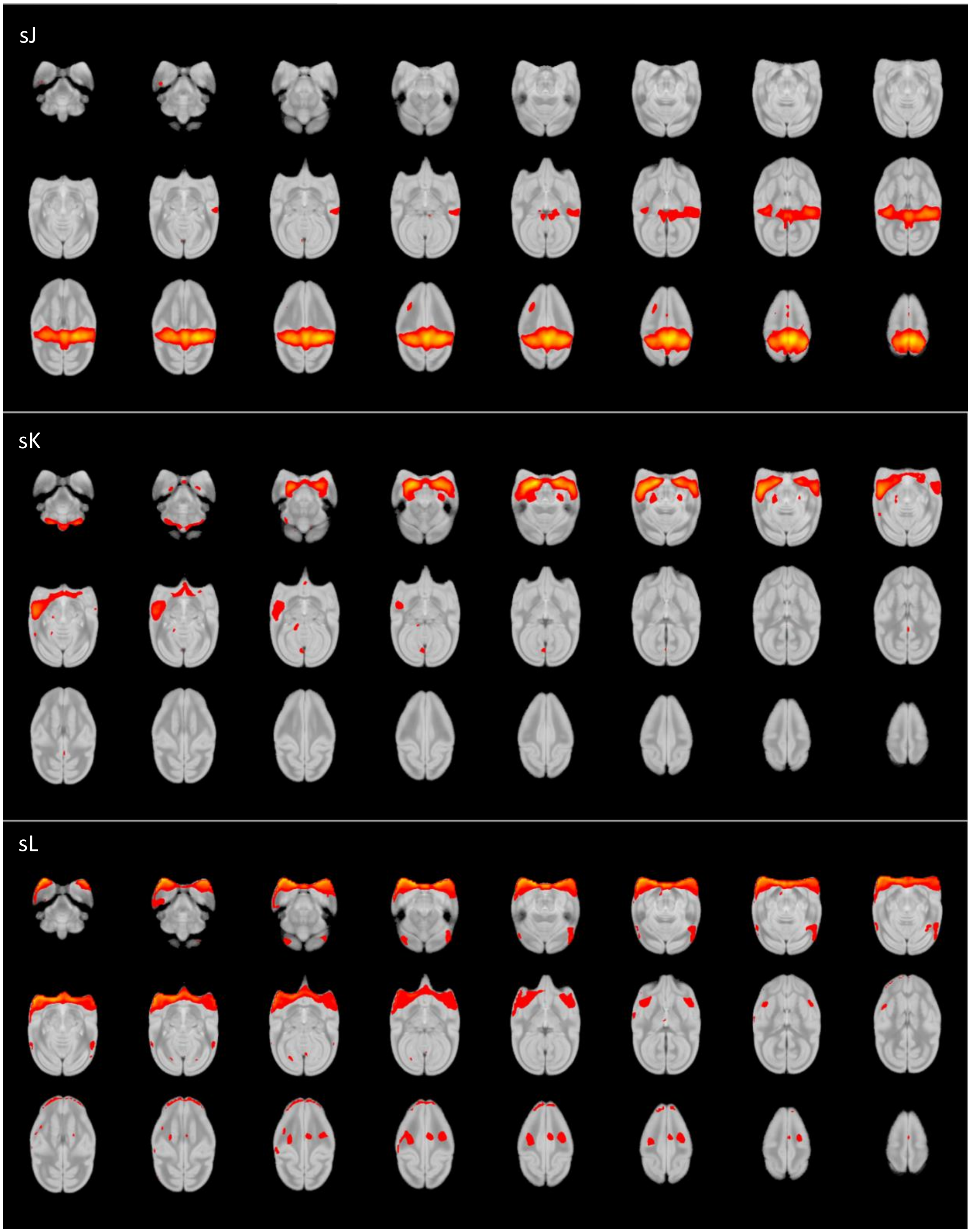
sJ
sK
sL

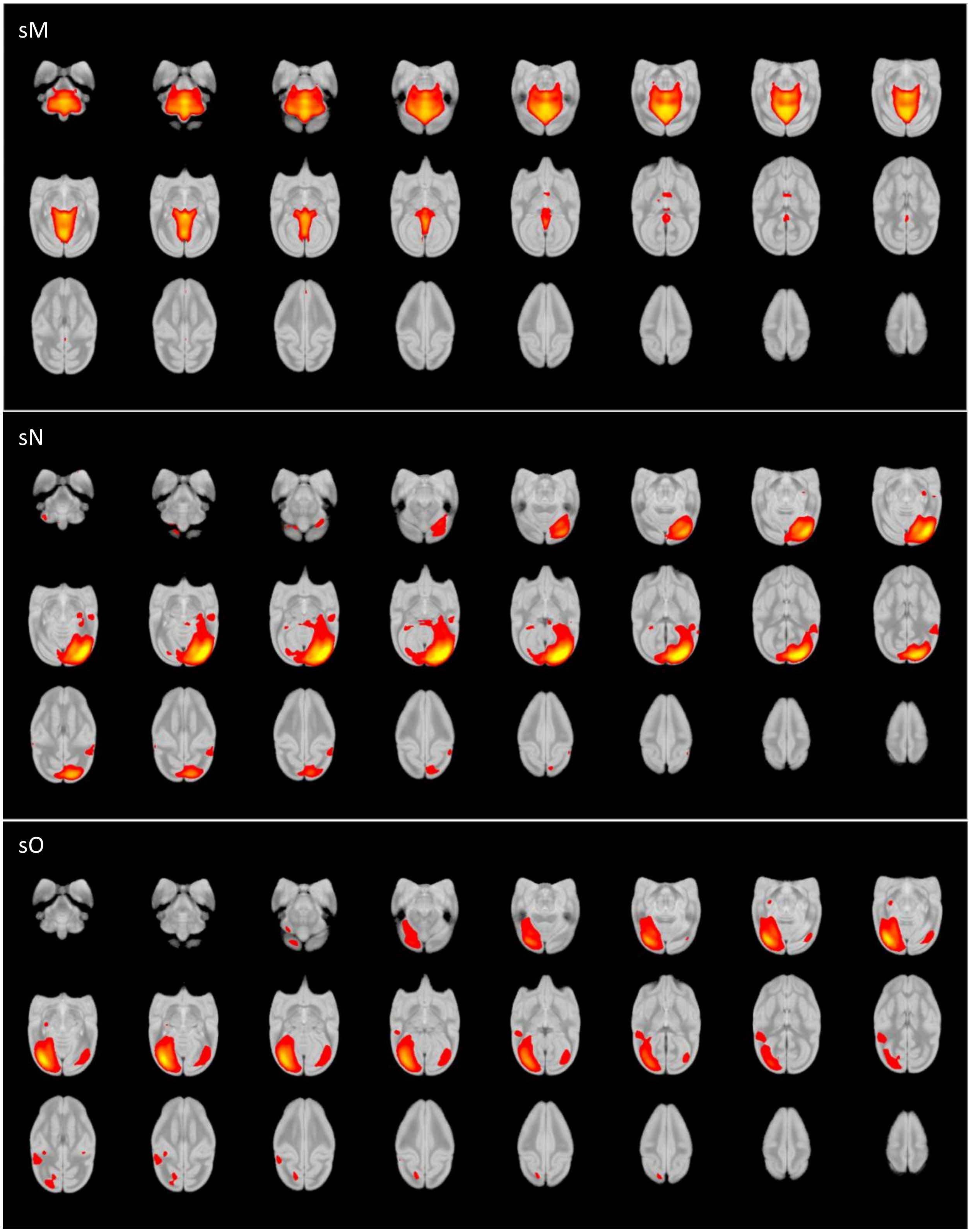
sM
sN
sO

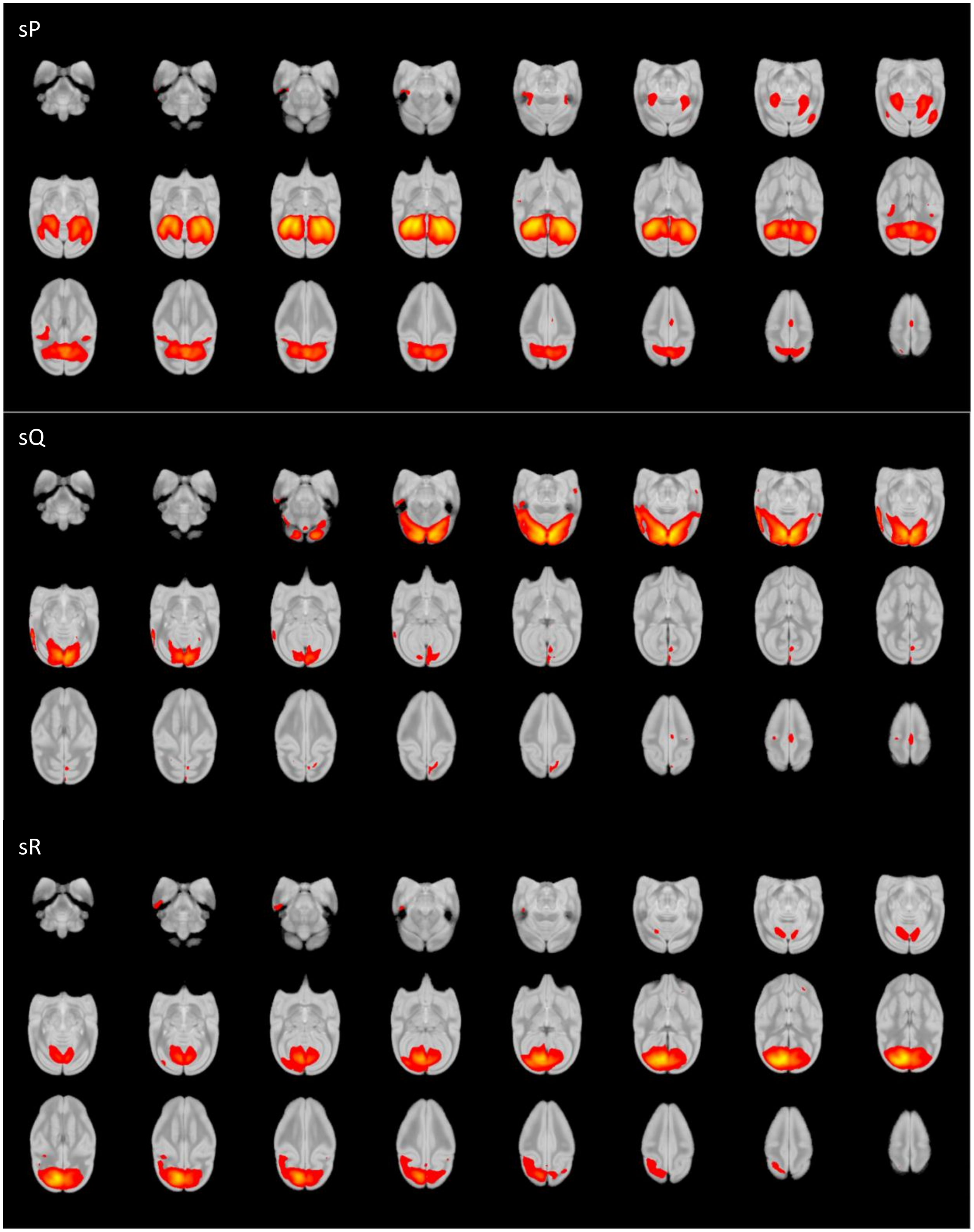
sP
sQ
sR

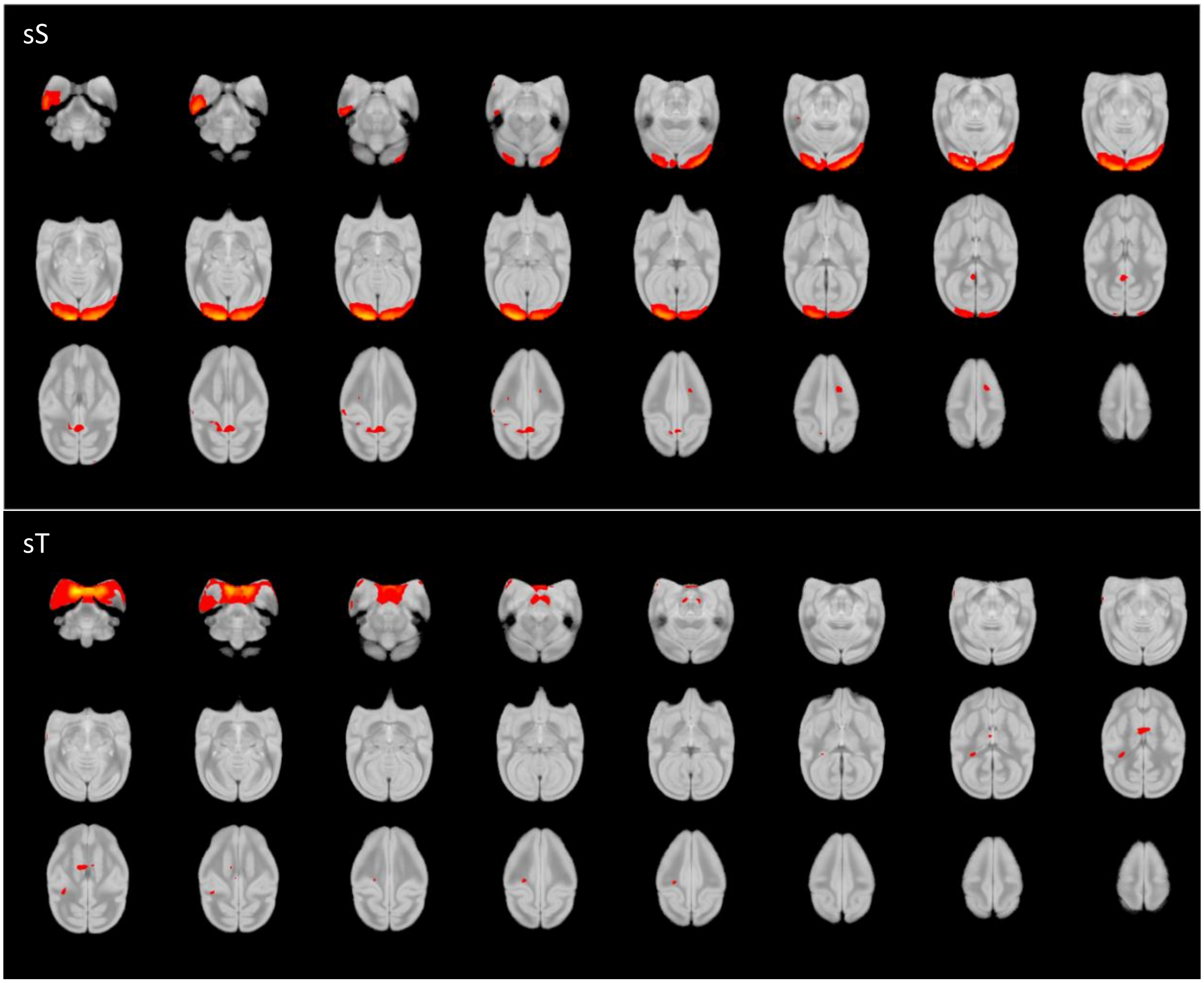


**Figure S2.**

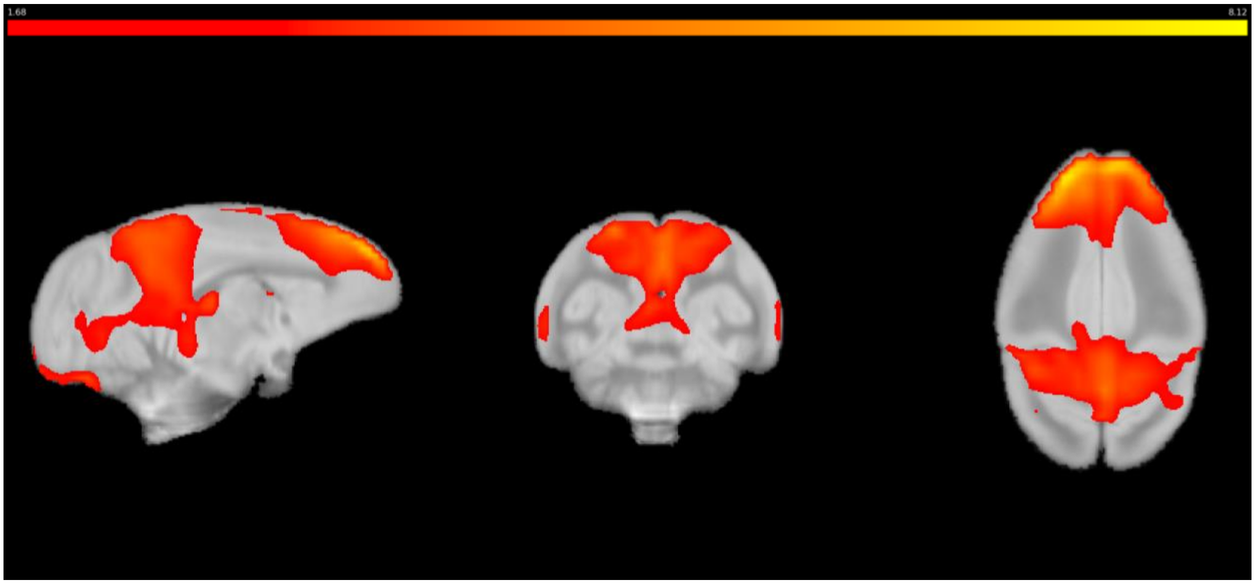


**Figure S3.**
